# Streptomycin and Pyridomycin drug molecules provide the most potential inhibitors against Rv3871 to cure Tuberculosis

**DOI:** 10.1101/2022.07.19.500019

**Authors:** Shikha Bhushan, Yashpal Singh Raja, Surya Mishra

**Affiliations:** INSTITUTE OF APPLIED MEDICINES AND RESEARCH DUHAI GHAZIABAD; SHARDA UNIVERSITY GREATER NOIDA; SHRI RAMSWAROOP MEMORIAL UNIVERSITY LUCKNOW

**Keywords:** Tuberculosis, Drug target, Molecular Docking, Screening of antituberculosis drug

## Abstract

The ESX-1 secretion system of Mycobacterium tuberculosis delivers bacterial virulence factors to host cells during infection. The most abundant factor, the ESAT-6/CFP-10 dimer, is targeted for secretion via a C-terminal signal sequence on CFP-10 that is recognized by the cytosolic ATPase, Rv3871. ATPase that is a component of the ESAT-6/CFP-10 secretion system. ESX locus contains genes encoding conserved secretion machinery components termed EccCb1. these core components are required for ESAT-6/CFP-10 secretion [70, 71]. Rv3871 is a cytoplasmic protein connected with the Rv3870 and encoded by the EccCb1 gene. These proteins supply energy for the secretion process. Each of these ATPases is involved in targeting protein for ESX-1 secretion. EccCb1 binds a seven amino acid C-terminal signal peptide of CFT-10, which is required for secretion of the ESAT-6/CFP-10 complex. In this we target the Rv3871 seven amino acid c-terminal region and block it through multiple drugs so, it cannot be activated by the ATPases and not to supply energy for the secretion protein during infection (Tuberculosis). In the present study we have comparatively studied different Rv3871 inhibitory antagonist drug molecule which could be a potential drug molecule to inhibit ATPases on the Rv3871 molecule that is responsible for the energy supply. We have 16 antagonistic drug molecules as shortlisted based on previous studies, that block the target site on the Rv3871 molecule.

## Introduction

Mycobacterium tuberculosis (M. tuberculosis) is a Gram-positive bacterium that causes Tuberculosis (TB). TB is one of the oldest known diseases and is still is one of the major causes of mortality. Two million people die each year from TB worldwide. TB is primarily a pulmonary disease caused by the deposition of M. tuberculosis contained in aerosol droplets on the lung alveolar surfaces (Rachman, H., Strong, M., Ulrichs, T., Grode, L., Schuchhardt, J., Mollen Kopf, H., Kosmiadi, G.A., Eisenberg, D. and Kaufmann, S.H., 2006). M. tuberculosis host cell infections critically depends on the release of virulence factors through the specialized secretion system ESX-1 which belongs to the type VII secretion systems (Abdallah, A.M., Van Pittius, N.C.G., Champion, P.A.D., Cox, J., Lui rink, J., Vandenbroucke-Grauls, C.M., Appel Melk, B.J. and Bitter, W., 2007). ESX-1 includes different cytoplasmic and membrane bound proteins which together leads to the transportation of virulence factors from the pathogen to the host cell. The ESX-1 secretion system includes Rv3868, Rv3870, Rv3871 and Rv3877 transporter as its major components; EspC, ESAT6 and CFP10 are the virulence small proteins secreted by ESX-1 system (Carlsson, F., Joshi, S.A., Rangell, L. and Brown, E.J., 2009). The objective of current proposal is to determine the Screening of drug molecules against Rv3871 transporter of Mycobacterium tuberculosis ESX-1 system using Auto dock, Auto dock vina for Molecular Docking and PyMol for comparing results which obtained from run by Command prompt. The Rv3871 enzyme binds to ESAT6/CFP10 and is involved in translocation of virulence factors across the bacterial cell wall (Fortune, S.M., Jaeger, A., Sar racino, D.A., Chase, M.R., Sassetti, C.M., Sherman, D.R., Bloom, B.R. and Rubin, E.J., 2005). The Rv3870+Rv3871 complex binds to ESAT6/CFP10 virulence factor and translocate across the bacterial cell wall. The Rv3877 transporter is located on the cytoplasmic membrane and facilitates the translocation of ESX-1 virulence proteins across the membrane. Screening of drug molecules against the Rv3871 will help to block the Target sites, and understand the virulence factor secretion by ESX-1 system. Structural basis of ESAT6/CFP10 recognition by Rv3871 enzyme will provide the platform for designing specific inhibitors (Xu, J., Laine, O., Masciocchi, M., Manoranjan, J., Smith, J., Du, S.J., Edwards, N., Zhu, X., Fenselau, C. and Gao, L.Y., 2007), which may block the Rv3871 secretion by ESX-1 system. The Screening of drug molecules against Rv3871 structure might reveal the Target site around Rv3871 and block the ATP site which provide the supply of energy to activate the Rv3871 molecule that cause the virulency during infection (Chen, J.M., Pojer, F., Blasco, B. and Cole, S.T., 2010). The Rv3871 transporter from ESX-1 system is involved in translocation of the virulence proteins across the cytoplasmic membrane. It is a cytoplasmic membrane protein that comprises 591 residues. The Screening of drug molecules against Rv3871 will explain it ATPases binding sites on the Rv3871 to activated it that for energy supply of translocation of virulence proteins across the membrane. A detailed understanding of the virulence protein secretion mechanism by ESX-1 system will be critical for *M. tuberculosis* drug development (Chen, J.M., Pojer, F., Blasco, B. and Cole, S.T., 2010). The Rv3871 transporter proteins constitute a pathway for virulence protein secretion by *M. tuebrculosis,* which can be targeted selectively for drug development (Bao YG, Qi ZF, Bao L. Nan Fang Yi, Ke Da Xue, Xue Bao 2009).

## Material and method

### Identification of drug target

The ESX-1 secretion system of *M. tuberculosis* delivers bacterial virulence factors into host cell during infection. The most abundant factors ESAT-6/CFP-10 dimer is targeted by ESX-1 system for secretion via C-terminal signal sequence of CFP-10, which recognize cytosolic AAA ATPase Rv3871(Di Giuseppe Champion, P.A., Champion, M.M., Manzanillo, P. and Cox, J.S., 2009). The EccCb1 (ESX-1 substrate protein Appeases, *Rv3871*) encoded outside the EccCb1 region is a target of cellular immunity (Millington, K.A., et al. 2011). In the patients with active and latent TB infection, the cellular immune response of EccCb1 is highly immunodominant as ESAT-6 and CFP-10(Sidders, Ben, et al. 2008; Millington, K.A., et al. 2011). Biochemical analysis has indicated that ESX-1 secreted substrates and AAA ATPase together form multi-protein complexes inside the cell (Di Giuseppe Champion, P.A., Champion, M.M., Manzanillo, P. and Cox, J.S., 2009). Both Rv3870/Rv3871 genes of ESX-1 system are essential for secretion of virulence factor ESAT6/CFP10(Fortune, S.M., Jaeger, A., Sar racino, D.A., Chase, M.R., Sassetti, C.M., Sherman, D.R., Bloom, B.R. and Rubin, E.J., 2005). The ATPase binding site on Rv3871+ EccCb1 transporter will reveal the overall blocking site of virulence factor secretion pathway adopted by *M. tuberculosis* ESX-1 system (Chen, J.M., Pojer, F., Blasco, B. and Cole, S.T., 2010). Screening analysis of these proteins will aid in designing novel inhibitors/drugs, which may block the ESX-1 secretion pathway in *M. tuberculosis.* (Feltcher, M.E., Sullivan, J.T. and Braunstein, M., 2010).

**Fig:1.**
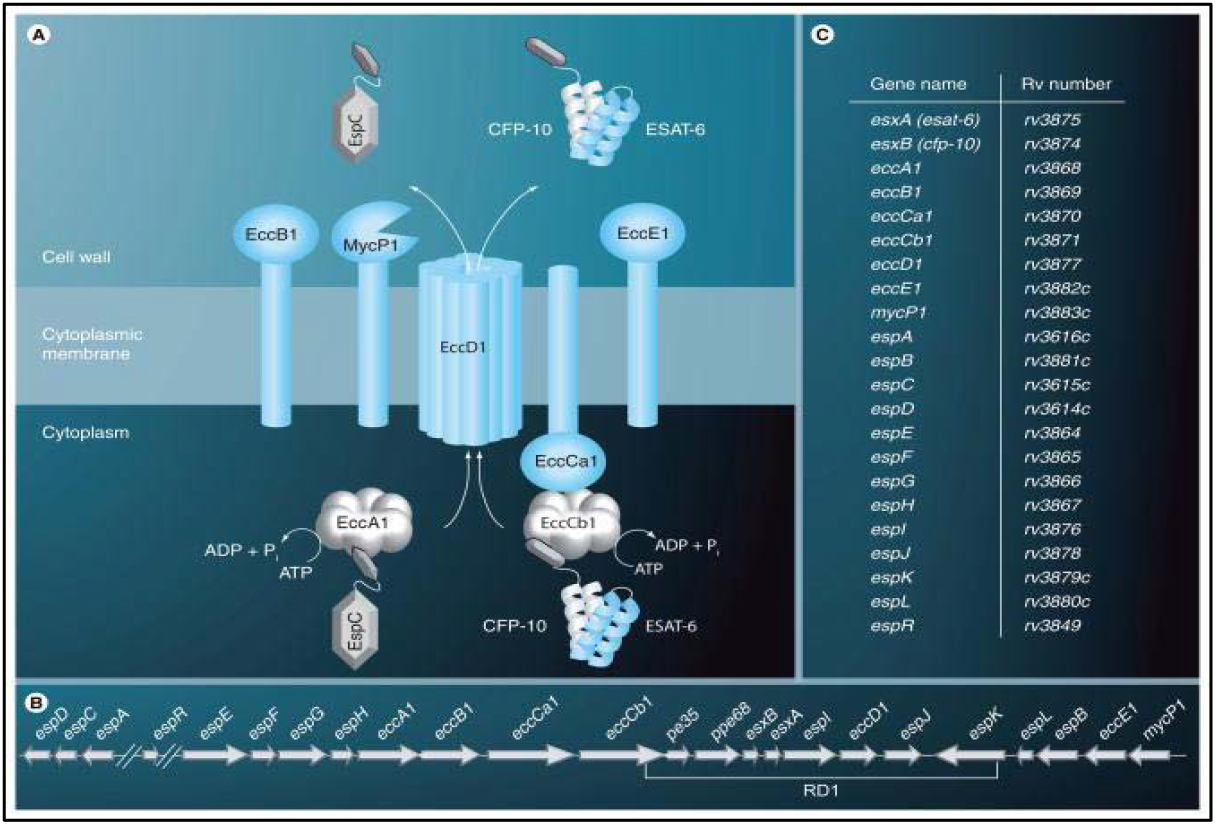
diagram of cytoplasmic membrane proteins and integral membrane proteins. The figure is adopted from Feltcher et al., 2010.

### Find Ligand for drug repurposing

**Table:1.** (Drug table)

| S. No | Candidate Drug molecules & Rv3871 | Compound ID (CID) PubChem |
| --- | --- | --- |
| Substrate | Rv3871 |  |
| 1 | Pyridomycin | Conformer3D_CID_3037036 |
| 2 | Streptomycin | Conformer3D_CID_19649 |
| 3 | Bed aquiline | Conformer3D_CID_5388906 |
| 4 | linezolid | Conformer3D_CID_441401 |
| 5 | Delamanid | Conformer3D_CID_6480466 |
| 6 | pA824 | Conformer3D_CID_456199 |
| 7 | Amikacin | Conformer3D_CID_37768 |
| 8 | Levofloxacin | Conformer3D_CID_149096 |
| 9 | Fluroquinolone | Conformer3D_CID_1526055 |
| 10 | Terizidone | Conformer3D_CID_65720 |
| 11 | Isoniazid | Conformer3D_CID_3767 |
| 12 | Pyrazinoic acid | Conformer3D_CID_1047 |
| 13 | Ofloxacin | Conformer3D_CID_4583 |
| 14 | Kanamycin | Conformer3D_CID_6032 |
| 15 | Clofazimin | Conformer3D_CID_2794 |
| 16 | Ethambutol | Conformer3D_CID_14052 |

#### Molecular Docking analysis

In this we used the MGL docking tool for the docking analysis. In this the grid box had formed that helped in the cover the target site. Retrieval of proteins sequence of Rv3871 Uniprot/PDB. Protein sequence information from database will be used for Clustal Omega EBI search for evolutionary Phylogeny analysis. Secondary structure will be determined using I-TASSER and structure inspection by using Pymol. Preparation of library of potential repurposing inhibitors for Tuberculosis (Rv3871). Preparation of 3D structure library of potential repurposing inhibitors for Tuberculosis (Rv3871) from PubChem. Molecular interaction analysis of individual drug inhibitor candidate against Rv3871 by Auto dock Vina, Auto Dock Tool (ADT) and PyMol.

#### Screening of Antagonistic drug against Tuberculosis

In the present study we have comparatively studied different Rv3871 inhibitory antagonist drug molecule which could be a potential drug molecule to inhibit ATPases on the Rv3871 molecule that is responsible for the energy supply. We have 16 antagonistic drug molecules as shortlisted based on previous studies, that block the target site on the Rv3871 molecule.

We selected the anti-tuberculosis drug that is Pyridomycin, streptomycin, Linezolid Bedaquiline, Delamanid, because these drugs block the ATPase site on the Rv3871 and stop the energy supply during infection that is why the proteins CFP-10/ESAT-6 do not pass the membrane. The Rv3871 that is cytoplasmic membrane protein that connect with the Rv3870 and EccCb1 together form a complex and it get activated by the binding of ATPase.

## Result

### Streptomycin and Pyridomycin are the most potential drug against Tuberculosis

The binding energies of these two drugs are comparatively lowest-6.5 and −7.4 kcal/mol respectively. These drugs cover the almost amino acids that is present on the Rv3871 loop and block them active sites loop (377AAKSGKTT384) and (574APY576) of the Rv3871 molecules.

**Fig:2.**
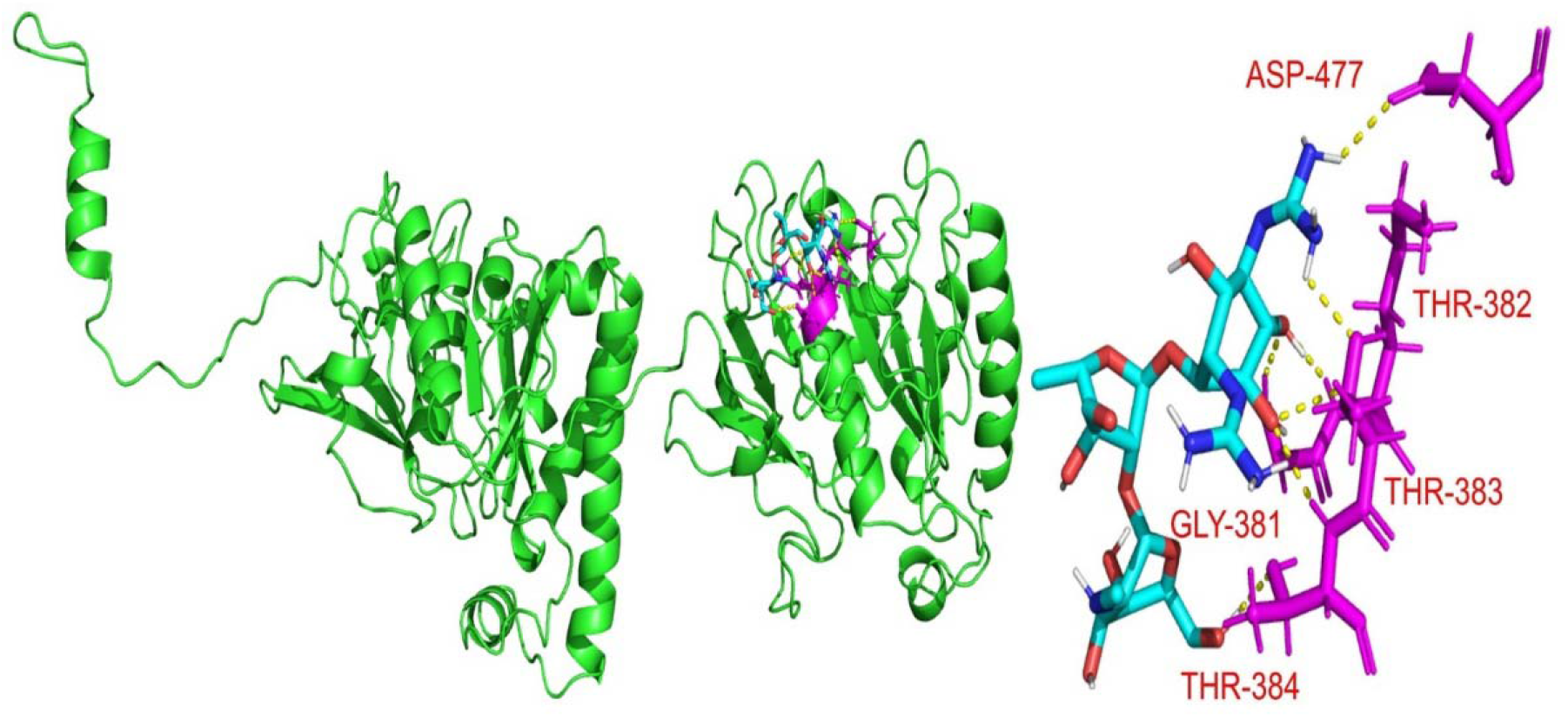
In this result the Streptomycin block the target site (that is bind with amino acid is ASP-477, THR-382, THR-383, GLY-381) these all the amino acid present on the target site.

**Fig:3.**
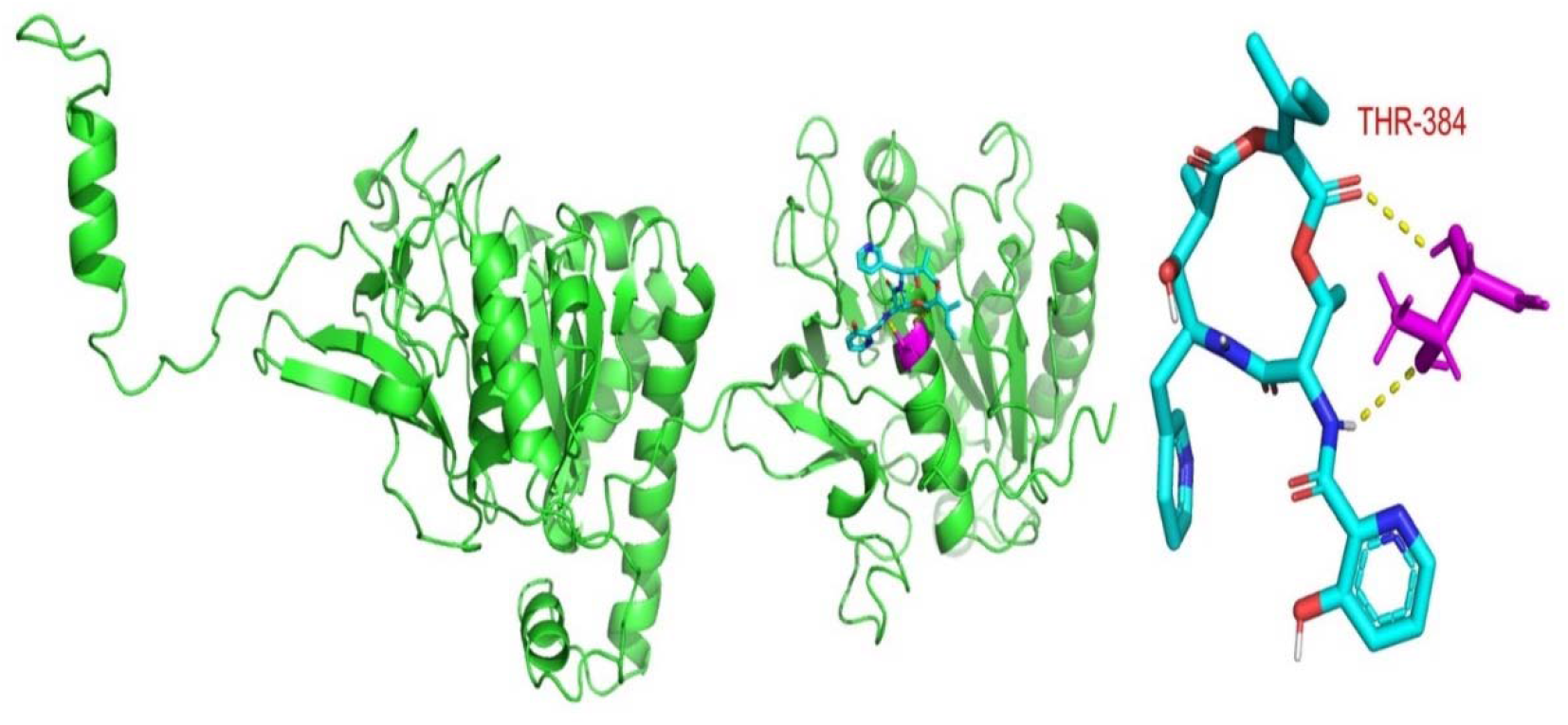
In this result the Pyridomycin block the target site (that is bind with amino acid is THR-384) these all the amino acid present on the target site.

**Fig: 4.**
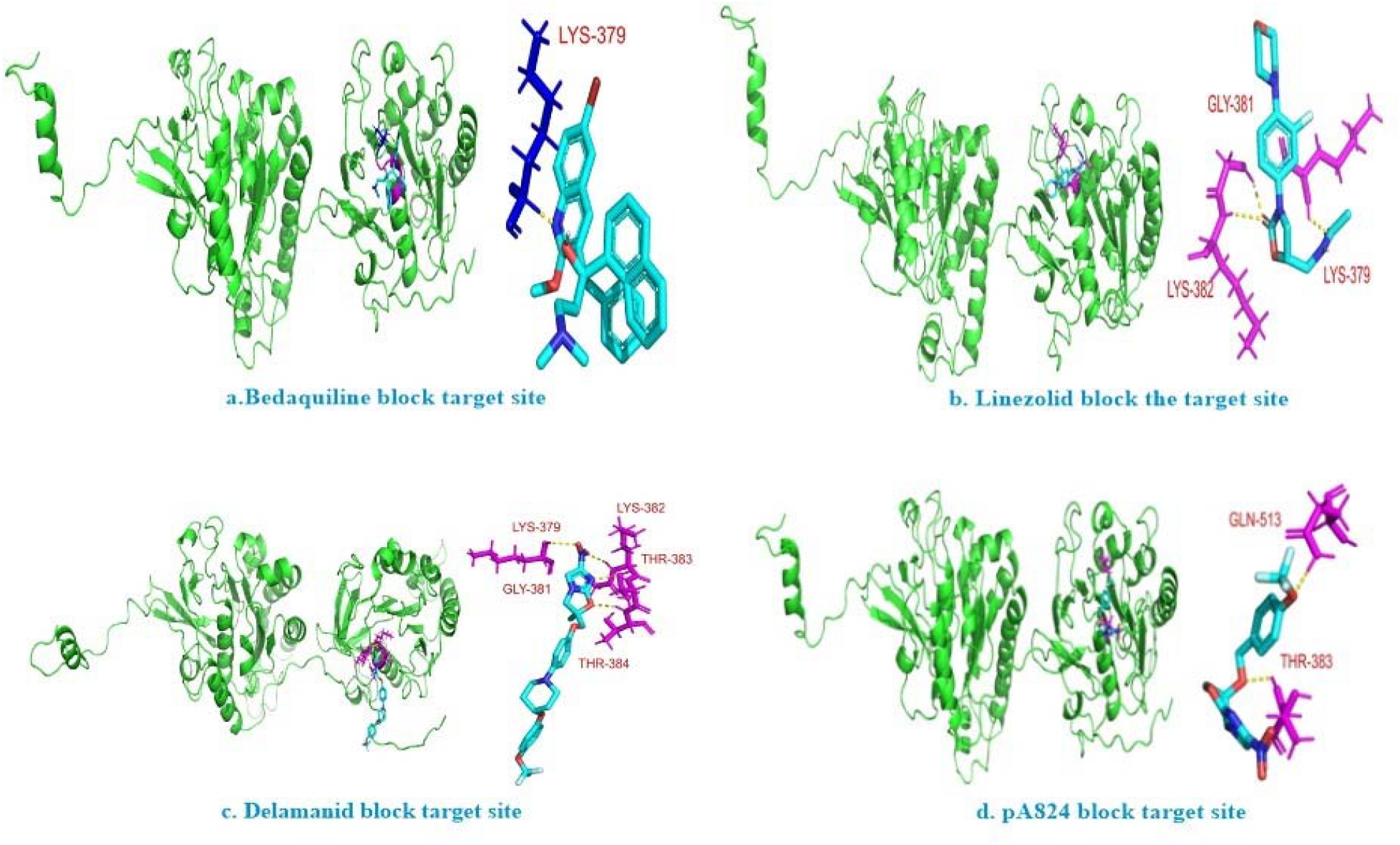
In the dig, a. Bedaquiline bind with the target amino acidLYS-379. b. Linezolid bind with the target amino acid GLY-381, LYS-379, LYS382. c. Delamanid bind with the LYS-382, LYS-379, THR-383, THR-384, GLY-381 target amino acid. d.pA824 bind with target amino acid the THR-383, and a non target amino acid GLN-513.

**Fig: 5.**
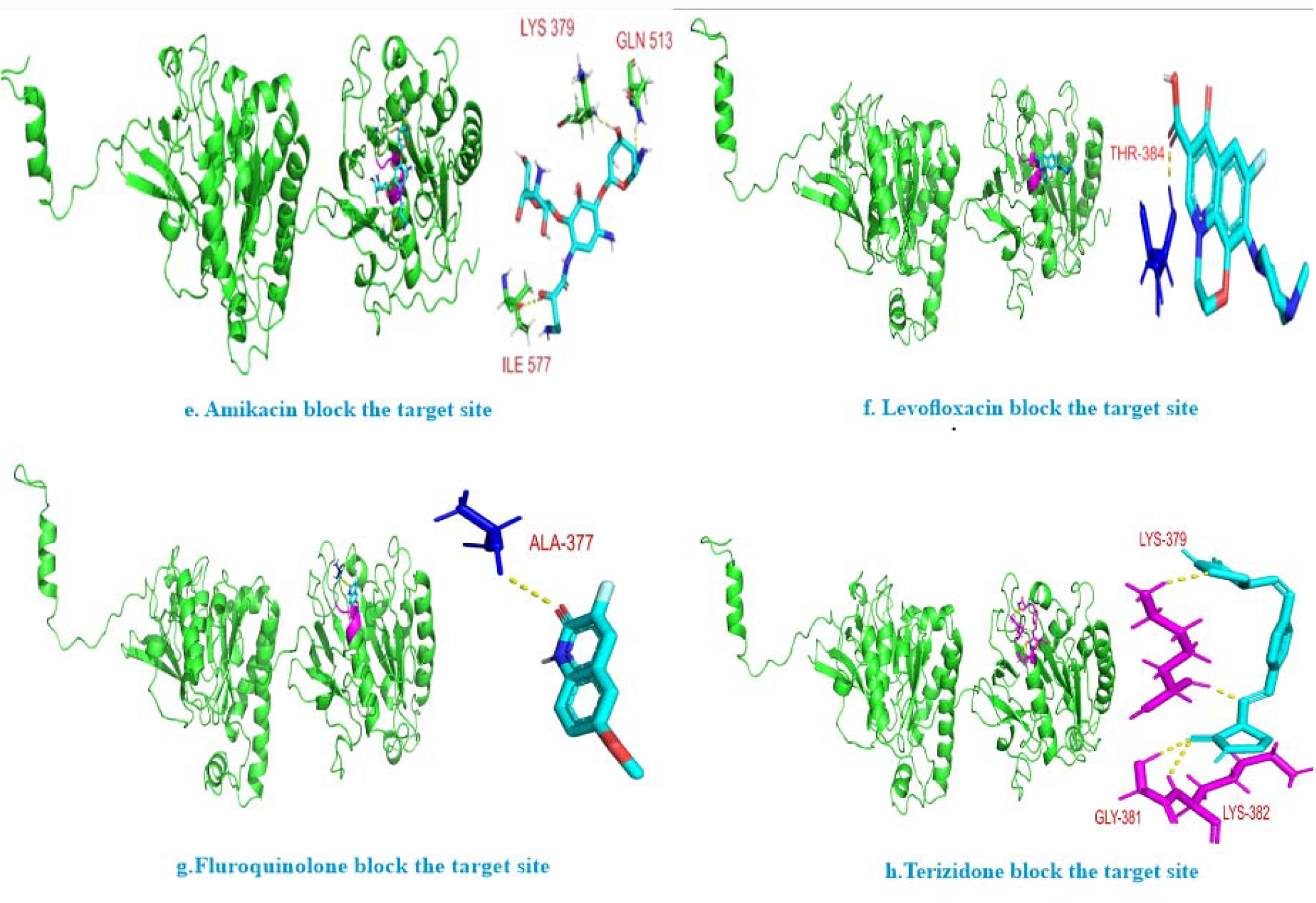
In the dig, e. Amikacin binds with amino acid LYS-379, and two non-target is ILE-577, GLN-513. f. Levofloxacin bind with the THR-384 target amino acid. In the g. Fluroquinolones bind with the target amino acid ALA-377. h. Terizidone bind with the target amino acid LYS-379, GLY-381, LYS-382.

**Fig: 6.**
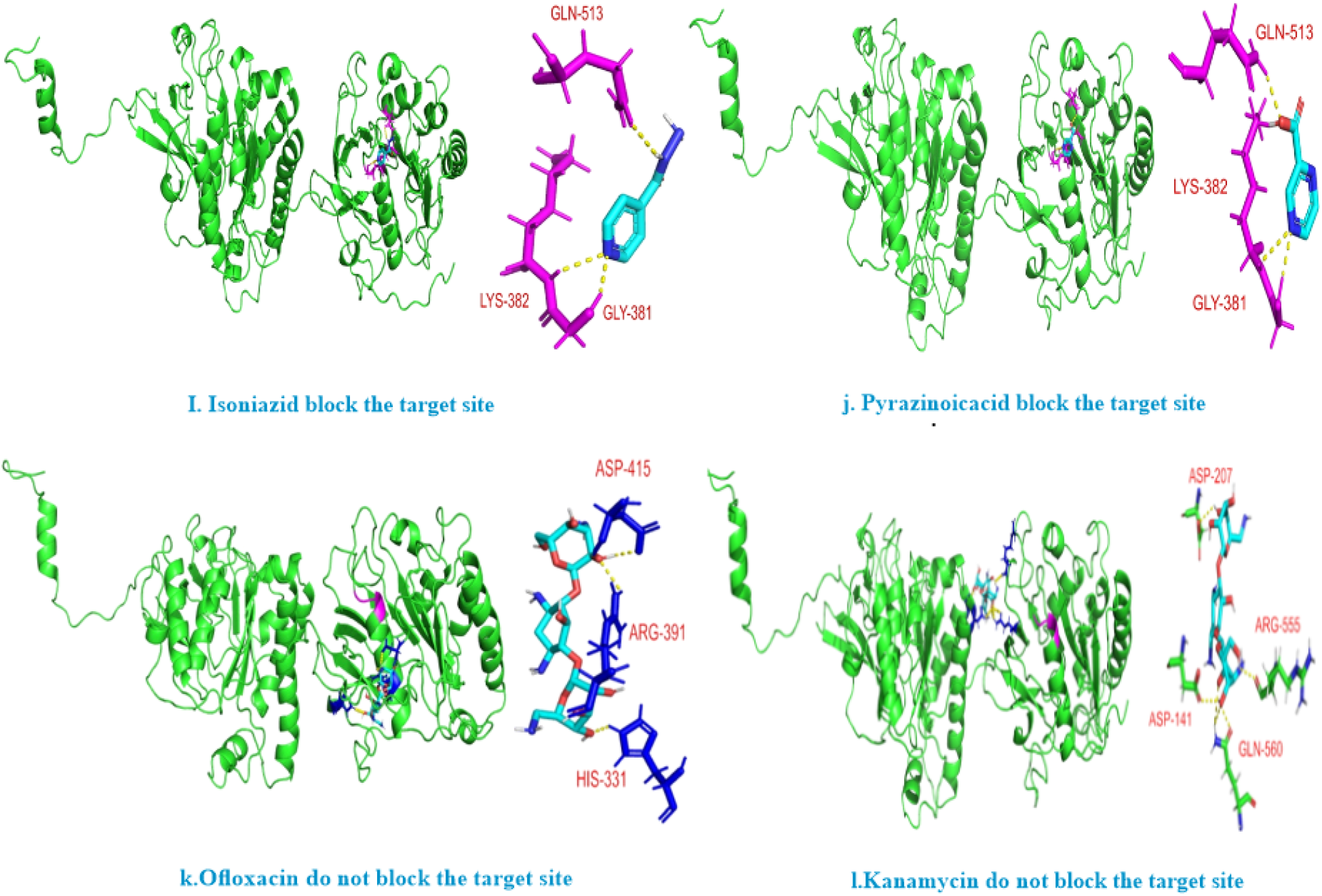
In. Isoniazid bind with the target amino acid GLY-381, LYS-382 and a non target site GLN-513. J. Pyrazinoic acid bind with the target amino acid GLY-381, LYS-382, and a non taget amino acid is GLN-513. K. Ofloxacin bind with the all non target amino acid are ASP-415, ARG-391, HIS-331. I. Kanamycin bind with all the non target site it don not block the target site like ASP-141, GLN-513, ASP-277, ARG-555.

**Table:2.** (Drug table blocking amino acid of active site with binding energy)

| S.No. | Drug Molecules | Binding Energy | Blocking amino acids of active site | Amino acids binding to ATP |
| --- | --- | --- | --- | --- |
| Substrate | ATP |  |  | ALA 377, ALA 378, SER 380, LYS 379, GLY 381, LYS 382, THR 384, THR 383, ILE 385, ALA 574, PRO 575, TYR 576 |
| 1 | Pyridomycin | -7.4 | THR-384 |  |
| 2 | Streptomycin | -6.5 | ASP-477, THR-382, THR-383, THR-384, GLY-381 |  |
| 3 | Bedaquiline | -6.3 | LYS-379 |  |
| 4 | linezolid | -6.2 | GLY-381, LYS-379, LYS-382 |  |
| 5 | Delamanid | -6.1 | LYS-379, LYS-382, GLY-381, THR-383, THR-384 |  |
| 6 | pA824 | -5.9 | GLN-513, THR-383 |  |
| 7 | Amikacin | -5.8 | LYS-379, GLN-513, ILE-577 |  |
| 8 | Levofloxacin | -5.8 | THR-384 |  |
| 9 | Fluroquinolone | -5.0 | ALA-377 |  |
| 10 | Terizidone | -4.8 | LYS-379, GLY-381, LYS-382 | ALA 377, ALA 378, SER 380, LYS 379, GLY 381, LYS 382, THR 384, THR383, ILE 385, ALA 574, PRO 575, TYR 576 |
| 11 | Isoniazid | -4.7 | GLN-513, LYS-382, GLY-381 |  |
| 12 | Pyrazinoicacid | -4.3 | GLN-513, LYS-382, GLY-381 |  |
| 13 | Ofloxacin | -6.4 | ASP-415, ARG-391, HIS-331 |  |
| 14 | Kanamycin | -5.8 | ASP-207, ARG-555, ASP141, GLN-560 |  |
| 15 | Clofazimin | -7.3 | GLY-381 |  |
| 16 | Ethambutol | -3.0 | ARG-391, ASP-415, ASP-491 |  |

## Conclusion

In the present study we have comparatively studied different Rv3871 inhibitory antagonist drug molecule which could be a potential drug molecule to inhibit ATPase on the Rv3871 molecule that is responsible for the energy supply. the Pyridomycin and Streptomycin molecules were observed to specifically bind to the key active site loop (377AAKSGKTT384) and (574APY576) of the Rv3871 molecules. The Binding energy of the Pyridomycin and Streptomycin was also comparatively lowest −7.4 and −6.5 kcal/mol respectively. Out of these two the Streptomycin drug is best inhibitory drug against Tuberculosis, because it blocks almost amino acid (ALA 377, ALA 378, SER 380, LYS 379, GLY 381, LYS 382, THR 384, THR383, ILE 385, ALA 574, PRO 575, TYR 576) on the target site. Hence, overall, from the present study we may conclude that the Streptomycin drug candidate would be a highly potential drug to inhibit ATPase on the Rv3871 and hence the drug could be a potential antituberculosis drug to inhibit the energy supply to the Rv3871 during infection. Although in vivo and human trials would be an essential obligation to the said conclusions, which may be addressed in future prospects of the present study.

